# Cancer metabolic subtypes and their association with molecular and clinical features

**DOI:** 10.1101/2021.12.16.472974

**Authors:** Enrico Moiso, Paolo Provero

## Abstract

Alterations of metabolic pathways in cancer have been investigated for many years, beginning way before the discovery of the role of oncogenes and tumor suppressors, and the last few years have witnessed a renewed interest in this topic. Large-scale molecular and clinical data on tens of thousands of samples allow us today to tackle the problem from a general point of view.

Here we show that transcriptomic profiles of tumors can be exploited to define metabolic cancer subtypes, which can be systematically investigated for association with other molecular and clinical data. We find thousands of significant associations between metabolic subtypes and molecular features such as somatic mutations, structural variants, epigenetic modifications, protein abundance and activation; and with clinical/phenotypic data including survival probability, tumor grade, and histological types.

Our work provides a methodological framework and a rich database of statistical associations, accessible from metaminer.unito.it, that will contribute to the understanding of the role of metabolic alterations in cancer and to the development of precision therapeutic strategies.

## Introduction

The last few years have witnessed a renewed interest in the metabolism of cancer, with the reprogramming of energy metabolism by cancer cells having been recently added, together with immune evasion, to the fundamental oncogenic processes collected under the heading of “Hallmarks of Cancer” [1].

Today, next-generation sequencing and other high-throughput techniques provide us with the unprecedented opportunity to take the study of the role of metabolic alterations in cancer progression to a new level, complementing the in-depth study of specific instances of metabolic changes with unbiased assessments of cancer metabolism and its relationship with the other hallmarks. In particular the TCGA project provides us with a systematic repository of genomic, epigenomic, transcriptomic, and clinical data on tens of thousands of samples of many tumor types. For example, a recent study [2] demonstrated the close relationship between patterns of copy-number alterations and metabolic phenotypes. Another study [3] showed the clinical relevance of metabolic pathways, as reflected by gene expression profiles of metabolic genes, to patient survival.

In this work we undertake a systematic analysis of the associations between metabolic profiles of human tumors and their molecular and clinical features. Our aim is to underline the role of metabolism in precision cancer medicine, by showing that metabolic profiling can classify tumors into classes that are significantly different in both their molecular-level features (somatic mutations, structural genomic variants, epigenetic modifications) and in their clinical aspects (survival probability, tumor grade, histological type). While intrinsically correlative in nature, this analysis can serve as a guide to mechanistic studies linking metabolism to other molecular features, and to the exploration of precision therapeutic approaches informed by tumor metabolism.

While, ideally, such a study should be based on direct assays of metabolite abundance in primary tumors, these are not yet available on a large scale. Therefore we use transcriptomic profiles of tens of thousands of samples of many tumor types as a proxy of metabolomic assays to classify them into metabolic subtypes. We then systematically correlate such subtypes with molecular and clinical data using appropriate statistical tests, thus generating a database of thousands of associations.

## Results

### Transcriptome-based classification of tumors into metabolic subtypes

For each tumor type with transcriptomic data available from the TCGA, and for each metabolic pathway defined as a set of genes by annotation databases such as KEGG [4,5], Reactome [6], and the Gene Ontology [7,8], we divided the patients into metabolic subtypes by clustering the samples using the expression profiles of the genes belonging to the pathway. Fig. 1 shows the KEGG pathways used in the analysis and the genes assigned to each pathway as a bipartite network.

**Figure 1:**
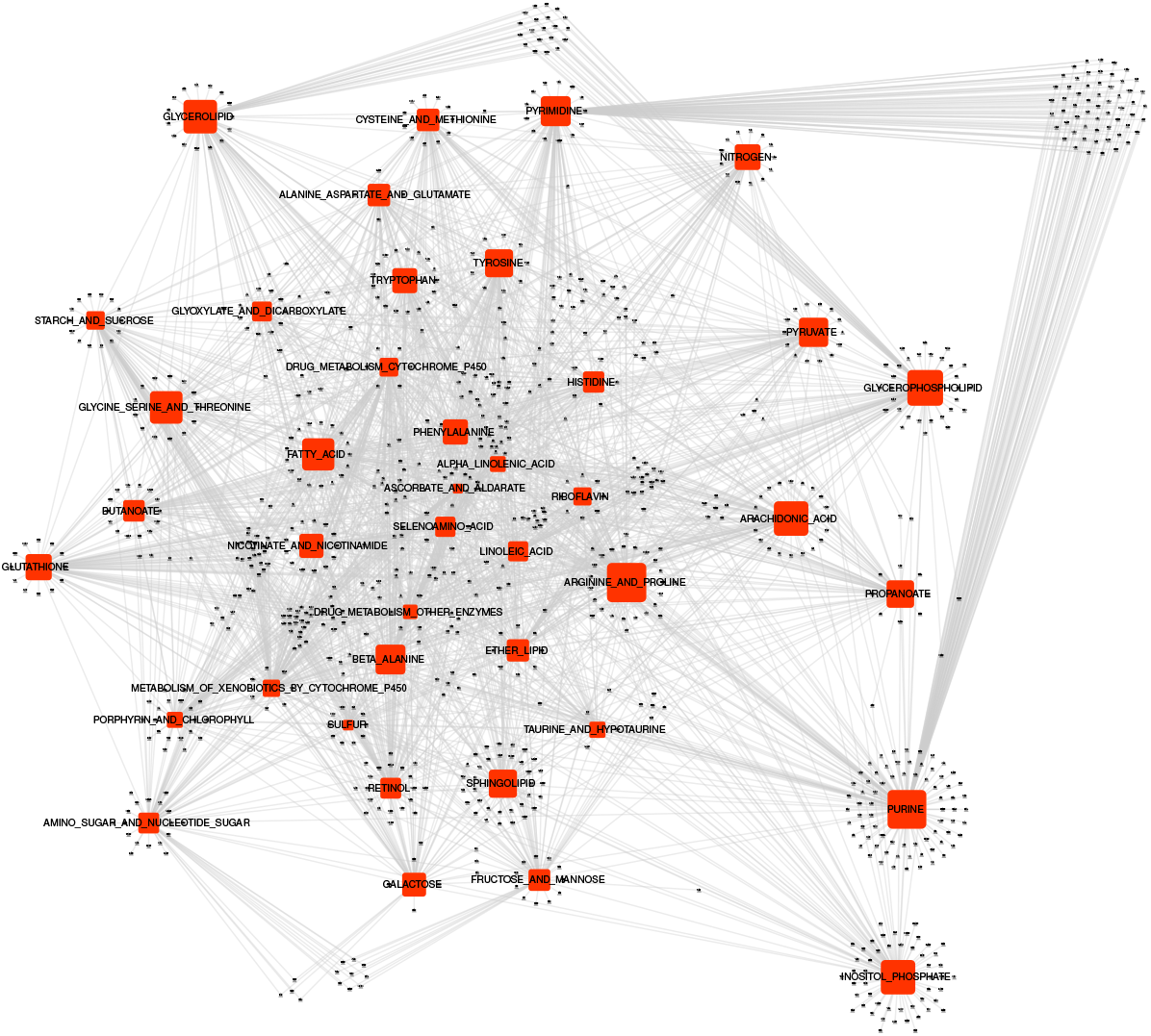
The KEGG pathways (red squares) used to cluster tumor samples and the respective genes (grey triangles) represented as a bipartite network. The size of the squares is proportional to the number of associated genes.

Compared to differential expression analyses used e.g. in [3], clustering by metabolic pathways correctly takes into account the fact that the genes associated to a pathway are often involved in either anabolic or catabolic processes. Therefore the activation of the pathway in a subset of patients is typically reflected by the transcriptional activation of the former and repression of the latter. For example, in Fig. 2, we show the clustering of low-grade glioma samples based on the “arachidonic acid metabolism” KEGG pathway: Each cluster is characterized by both up- and downregulated genes, a structure that would not be captured by simple differential expression of the pathway genes as a whole.

**Figure 2:**
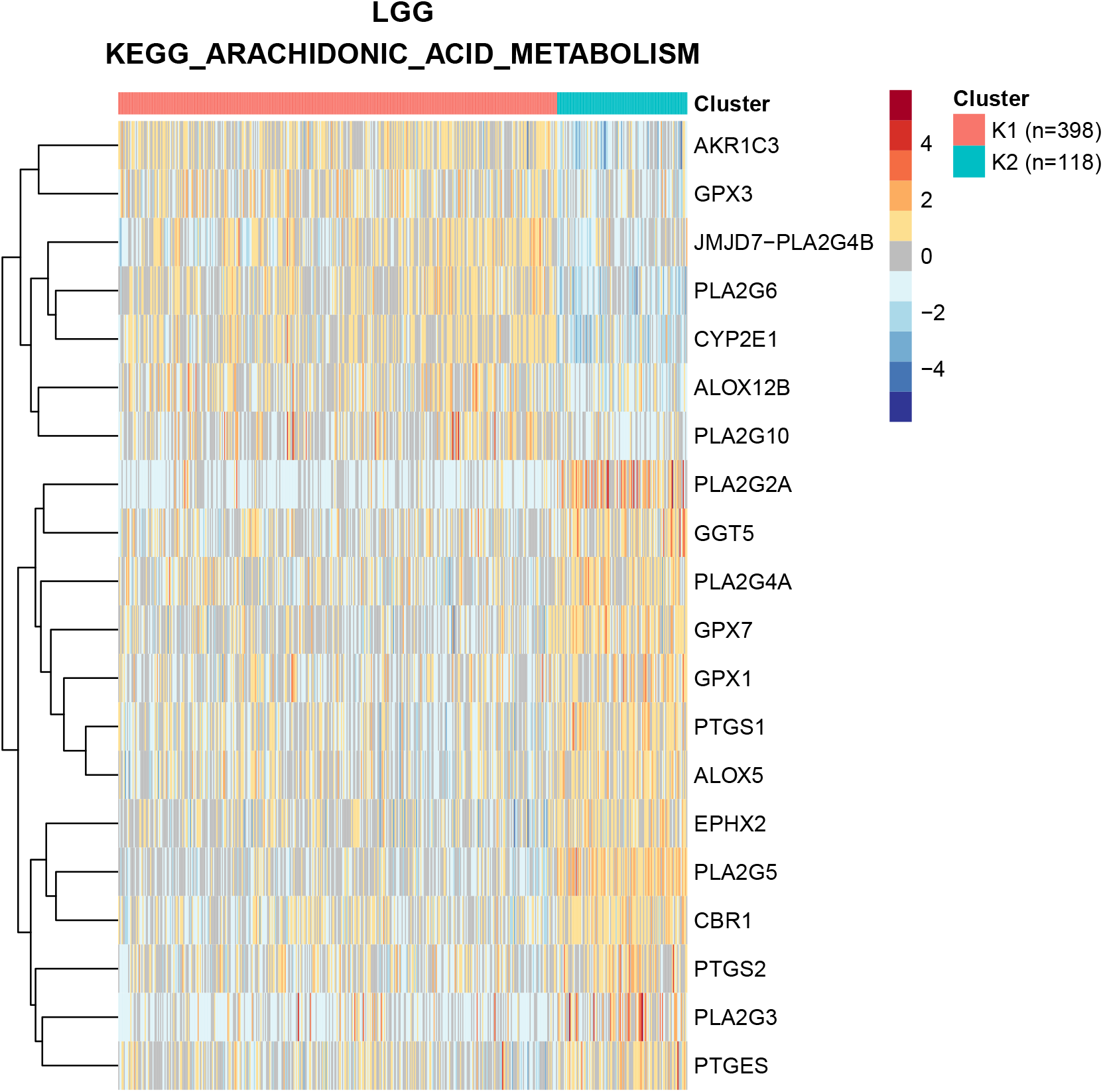
Clustering of low-grade glioma patients using the genes involved in arachidonic metabolism according to KEGG. Only the 20 genes most differentially expressed between the two clusters are shown.

To perform the clustering we used partitioning around medoids (PAM) [9], a more robust method compared to k-means [10] with respect to the presence of outliers. Cluster silhouette analysis allowed an unsupervised choice of the number of clusters. A total of 345 metabolic gene sets were used, as described in the Methods, on 38 tumor classes, thus performing a total of 13110 clusterings. In the majority of cases (9654, 73.6%) the samples were subdivided into *k* = 2 clusters, while in other cases we obtained up to 10 clusters. These clusters will be referred to as “metabolic subtypes.” The tumor types analyzed with the number of samples for which expression data were available are shown in Table 1.

**Table 1:** Number of samples used for each tumor class.

| ID | Tumor_type | Samples |
| --- | --- | --- |
| ACC | Adrenocortical Carcinoma | 79 |
| BLCA | Bladder Urothelial Carcinoma | 408 |
| BRCA | Breast Invasive Carcinoma | 1093 |
| CESC | Cervical Squamous Cell Carcinoma and<br>Endocervical Adenocarcinoma | 304 |
| CHOL | Cholangiocarcinoma | 36 |
| COAD | Colon Adenocarcinoma | 285 |
| COADREAD | (= COAD + READ) | 379 |
| DLBC | Lymphoid Neoplasm Diffuse Large B-cell<br>Lymphoma | 48 |
| ESCA | Esophageal Carcinoma | 184 |
| GBM | Glioblastoma Multiforme | 153 |
| GBMLGG | (= GBM + LGG) | 669 |
| HNSC | Head and Neck Squamous Cell Carcinoma | 520 |
| KICH | Kidney Chromophobe | 66 |
| KIPAN | (= KICH + KIRC + KIRP) | 889 |
| KIRC | Kidney Renal Clear Cell Carcinoma | 533 |
| KIRP | Kidney Renal Papillary Cell Carcinoma | 290 |
| LAML | Acute Myeloid Leukemia | 173 |
| LGG | Brain Lower Grade Glioma | 516 |
| LIHC | Liver Hepatocellular Carcinoma | 371 |
| LUAD | Lung Adenocarcinoma | 515 |
| LUNG | (= LUAD + LUSC) | 1016 |
| LUSC | Lung Squamous Cell Carcinoma | 501 |
| MESO | Mesothelioma | 87 |
| OV | Ovarian Serous Cystadenocarcinoma | 303 |
| PAAD | Pancreatic Adenocarcinoma | 178 |
| PCPG | Pheochromocytoma and Paraganglioma | 179 |
| PRAD | Prostate Adenocarcinoma | 497 |
| READ | Rectum Adenocarcinoma | 94 |
| SARC | Sarcoma | 259 |
| SKCM | Skin Cutaneous Melanoma | 103 |
| STAD | Stomach Adenocarcinoma | 415 |
| STES | (= STAD + ESCA) | 599 |
| TGCT | Testicular Germ Cell Tumors | 150 |
| THCA | Thyroid Carcinoma | 501 |
| THYM | Thymoma | 120 |
| UCEC | Uterine Corpus Endometrial Carcinoma | 176 |
| UCS | Uterine Carcinosarcoma | 57 |
| UVM | Uveal Melanoma | 80 |

**Table 2:** Molecular and phenotypic variables analyzed for association with metbolic subtypes

| class | description | type |
| --- | --- | --- |
| CNA | Copy number alterations | molecular |
| Methylation | Global DNA methylation | molecular |
| miRNA | microRNA expression | molecular |
| Mutations | Specific point mutations | molecular |
| Gene Mutations | Gene-level mutations | molecular |
| RPPA | Protein expression | molecular |
| Clinical | Clinical and histologic parameters | phenotypic |
| OS | Overall survival | phenotypic |
| RFS | Recurrence-free survival | phenotypic |

It is worth asking whether different metabolic gene sets cluster the patients of the same tumor in significantly different ways. To answer this question we computed the normalized mutual information between the cluster assignments obtained in each tumor with each metabolic gene set. The results are shown in Fig. 3, and suggest that, in most cases, different metabolic gene sets cluster tumor patients in clearly distinct ways.

**Figure 3:**
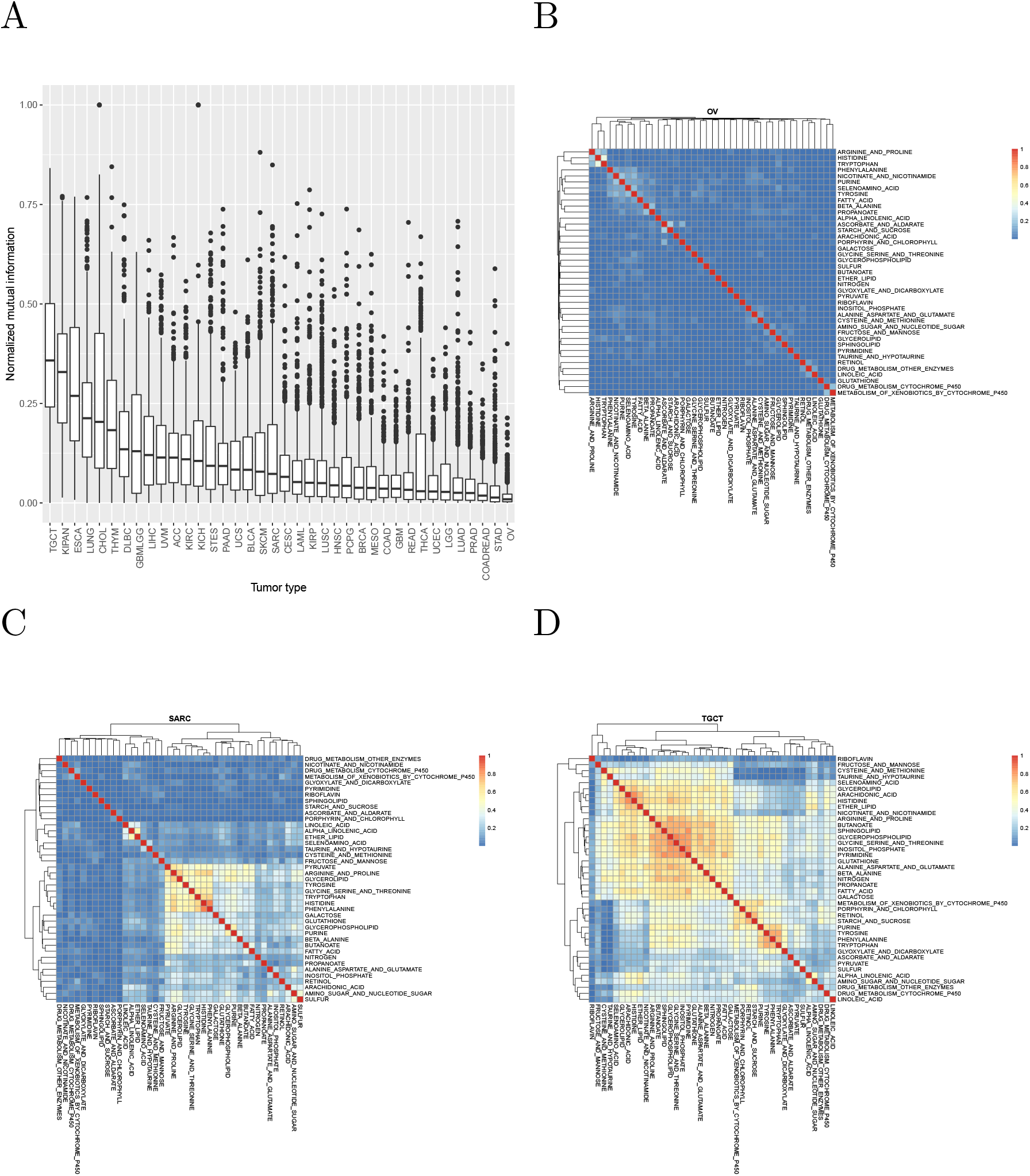
A: Normalized mutual information between clusterings of samples using KEGG metabolic gene sets. B, C, D: examples of tumor types with low, intermediate, and high mutual information.

### Associations between metabolic subtypes and molecular and clinical features

We then proceeded to systematically explore the statistical associations between metabolic subtypes and several molecular and clinical features of the corresponding tumors, classified into 9 variable classes. Statistical tests appropriate to the nature of the variables describing such features were chosen as described in the Methods. Bonferroni correction was applied separately to each variable type but to all tumor types together. Thus, for example, the Bonferroni correction for the association with miRNA expression takes into account all tests of association between all miRNAs and all metabolic gene sets across all tumor types. The molecular and phenotypic variables considered are shown in the following table:

In the following, to limit redundancy, we focus on the results obtained with the KEGG annotation database (a total of 99621 significant associations). Figure 4A shows the distribution of these associations among tumor types. Note that the number of significant associations for a given tumor type will depend in general on the number of available samples: indeed the Spearman correlation coefficient between number of associations and number of samples is 0.755. In Figure 4B we show the number of associations by KEGG metabolic pathway, suggesting a particular relevance of the metabolism of aminoacids and fatty acids in classifying tumors into classes characterized by different molecular and/or clinical characteristics.

**Figure 4:**
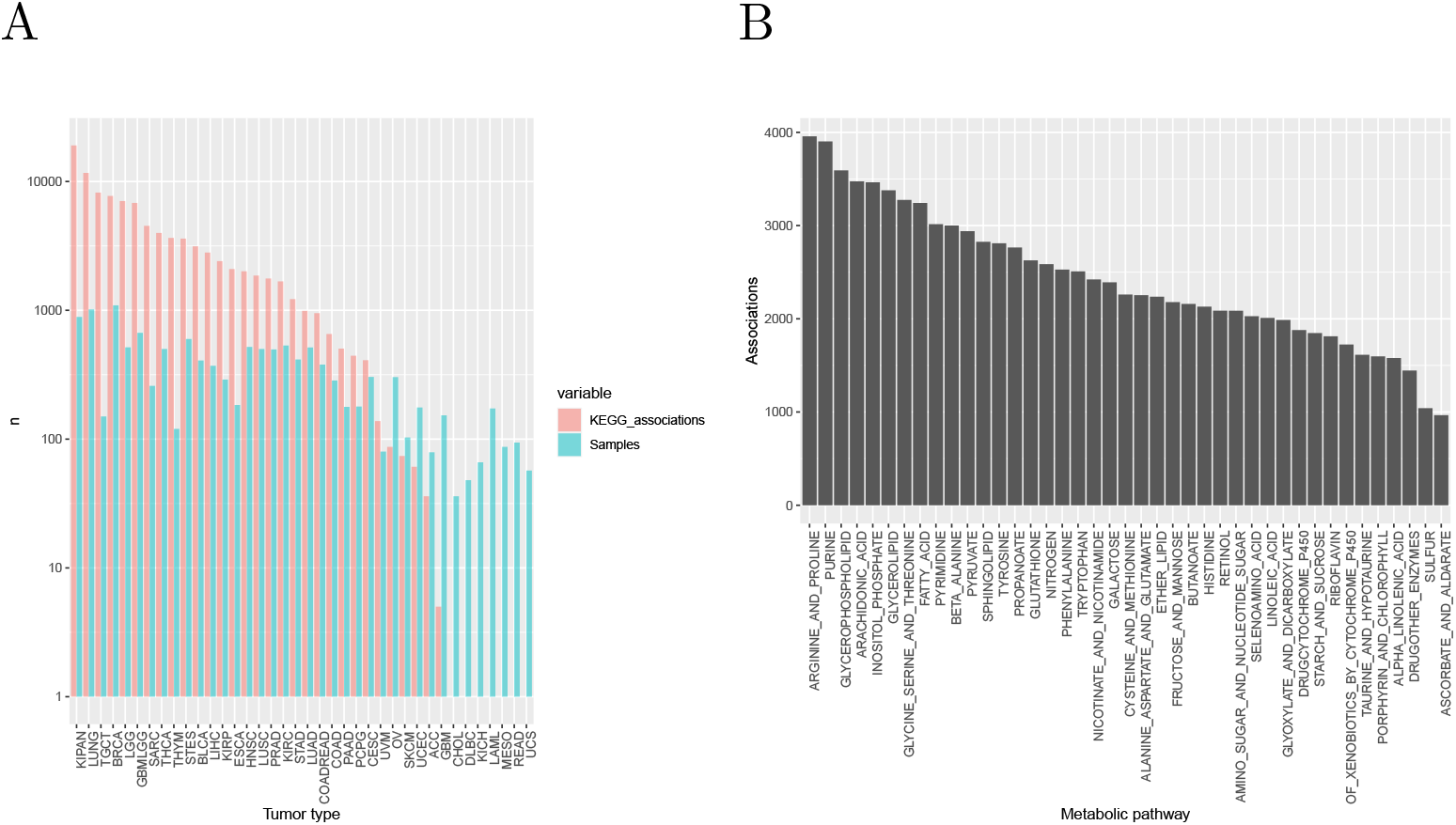
(A) Number of samples and of significant associations with molecular and clinical features by tumor type. (B) Number of significant associations by KEGG metabolic pathway.

### Recurrent associations

Of particular interest are recurrent associations, that is associations that are statistically significant in more than one tumor type (not considering the tumor types built from the union of smaller types, such as LUNG, KIPAN, etc.).

We found a total of 14937 such recurrent associations. Table 3 shows the most recurrent associations for each class of clinical and molecular variables. Notably, the only variable classes with no recurrent associations are point mutations: All the 45 associations with specific mutations and the 102 associations with gene-level mutations are specific of a single tumor type.

**Table 3:** Top recurrent associations for each class of variables

| Variable | Metabolism | Number of associations | Tumors associated |
| --- | --- | --- | --- |
| miRNA: hsa-mir-222 | INOSITOL<br>PHOSPHATE | 12 | BLCA, BRCA, CESC,<br>HNSC, KIRC, LGG, LIHC,<br>LUAD, SARC, STAD,<br>TGCT, THCA |
| Methylation | GLYCEROLIPID | 10 | BLCA, CESC, COAD,<br>HNSC, LGG, PCPG, SARC,<br>TGCT, THCA, THYM |
| Methylation | INOSITOL<br>PHOSPHATE | 10 | BLCA, COAD, KIRP, LGG,<br>LIHC, LUAD, PCPG,<br>TGCT, THCA, THYM |
| Methylation | PHENYLALANINE | 10 | BLCA, CESC, COAD, LGG,<br>LIHC, LUAD, OV, PCPG,<br>SARC, TGCT |
| Methylation | PURINE | 10 | BLCA, CESC, KIRP, LIHC,<br>LUAD, PCPG, SARC,<br>TGCT, THCA, THYM |
| Clinical: histological type | TYROSINE | 8 | BRCA, ESCA, LGG, PCPG,<br>SARC, THCA, THYM,<br>UCEC |
| RPPA: CCNB1\|Cyclin B1 | PURINE | 7 | BLCA, BRCA, KIRC, KIRP,<br>LUAD, TGCT, THYM |
| CNA: chr17p12 | FATTY ACID | 6 | COAD, KIRP, LGG, OV,<br>PAAD, TGCT |
| CNA: chr17q11 | GLYCEROLIPID | 6 | BRCA, LGG, LUSC, SARC,<br>TGCT, THYM |
| CNA: chr17q21 | GLYCEROLIPID | 6 | BRCA, HNSC, LGG, LUSC,<br>TGCT, THYM |
| CNA: chr3p | INOSITOL<br>PHOSPHATE | 6 | BLCA, BRCA, ESCA,<br>KIRC, LUSC, UVM |
| CNA: chr3p14 | INOSITOL<br>PHOSPHATE | 6 | BLCA, BRCA, ESCA,<br>KIRC, LUSC, UVM |
| CNA: chr3p21 | ARGININE AND<br>PROLINE | 6 | BRCA, ESCA, KIRC,<br>PRAD, SARC, THYM |
| CNA: chr3p21 | INOSITOL<br>PHOSPHATE | 6 | BLCA, BRCA, ESCA,<br>KIRC, LUSC, UVM |
| CNA: chr3q | PURINE | 6 | BRCA, ESCA, HNSC,<br>LUSC, UCEC, UVM |
| CNA: chr3q24 | PURINE | 6 | BRCA, ESCA, HNSC,<br>LUSC, UCEC, UVM |
| CNA: chr3q25 | PURINE | 6 | BRCA, ESCA, HNSC,<br>LUSC, UCEC, UVM |
| CNA: chr3q26 | PURINE | 6 | BRCA, ESCA, HNSC,<br>LUSC, UCEC, UVM |
| CNA: chr3q27 | PURINE | 6 | BRCA, ESCA, HNSC,<br>LUSC, UCEC, UVM |
| CNA: chr3q28 | FATTY ACID | 6 | BRCA, ESCA, HNSC, LGG,<br>LUAD, LUSC |
| CNA: chr3q28 | PURINE | 6 | BRCA, ESCA, HNSC,<br>LUSC, UCEC, UVM |
| CNA: chr3q29 | FATTY ACID | 6 | BRCA, ESCA, HNSC, LGG,<br>LUAD, LUSC |
| CNA: chr3q29 | PURINE | 6 | BRCA, ESCA, HNSC, LUSC, UCEC, UVM |
| OS | PROPANOATE | 2 | KIRC, LGG |
| OS | PURINE | 2 | ACC, KIRC |
| OS | PYRIMIDINE | 2 | ACC, LGG |
| OS | SPHINGOLIPID | 2 | KIRC, LGG |

Figure 5 shows the expression of microRNA miR-22 in the clusters of breast cancer and lung adenocarcinoma patients obtained with the genes associated to inositol phopsphate metabolism.

**Figure 5:**
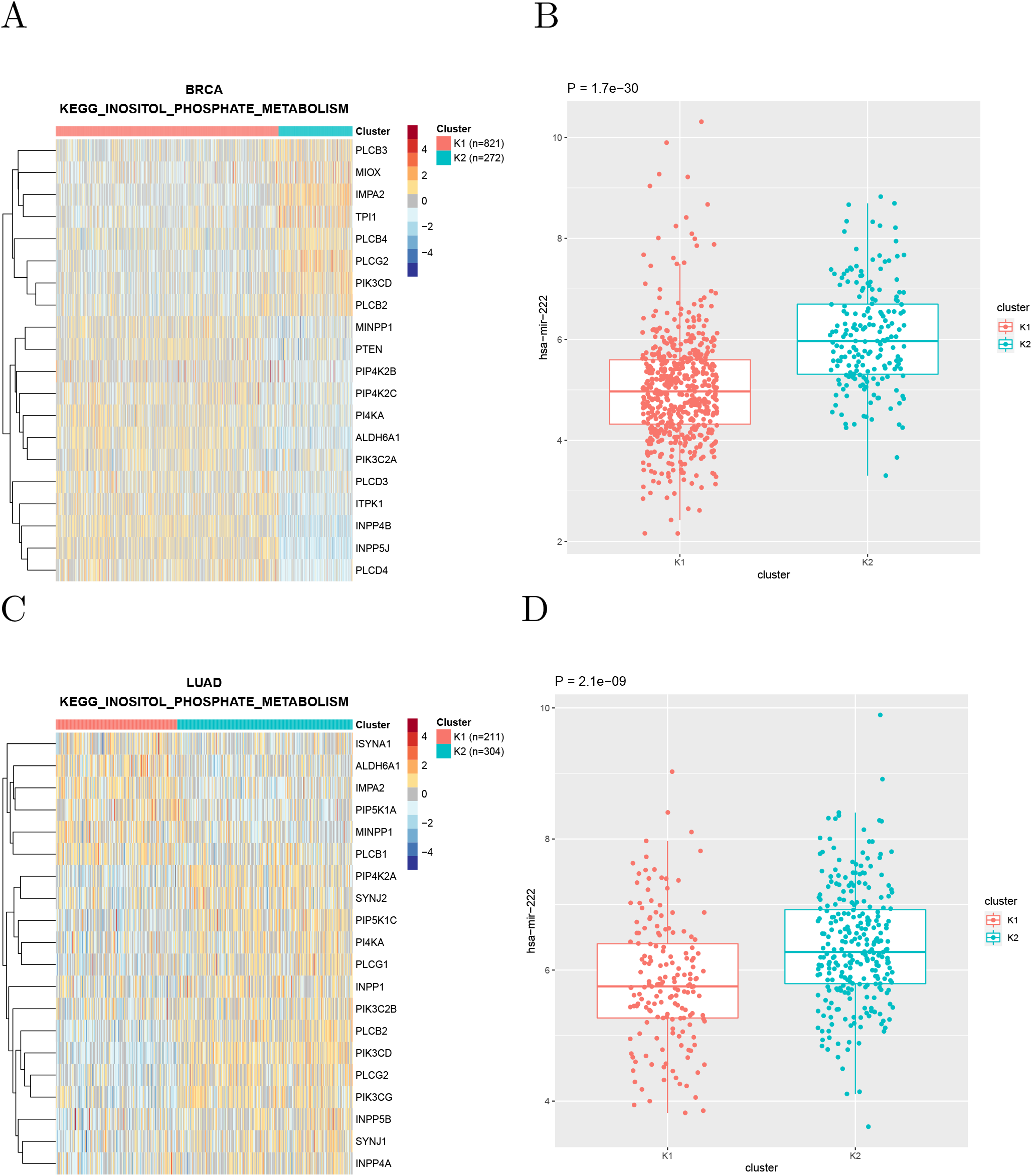
(A) Clustering of breast cancer samples using the genes associated to inositol phosphate metabolism according to KEGG (B) Expresson of miR-222 in the two clusters. (C, D) Same for lung adenocarcinoma samples

### Specific examples

In the following we show some examples of associations, some confirming known results and some suggesting new lines of investigation, especially in less widely studied tumor types.

### TP53 mutations and glucose metabolism

Mutant TP53 has been shown to promote the Warburg effect, possibly the most notoriuos metabolic hallmark of cancer, in breast and lung cancer cell lines [11]. Indeed clustering breast and lung tumors using the genes associated to glucose metabolism according to REACTOME shows, in both cancer types, the appearance of a cluster characterized by overexpression of several Warburg effect signature genes (such as GAPDH, PGK1, and PKM2) and by an enrichment in TP53 mutations (Fig. 6).

**Figure 6:**
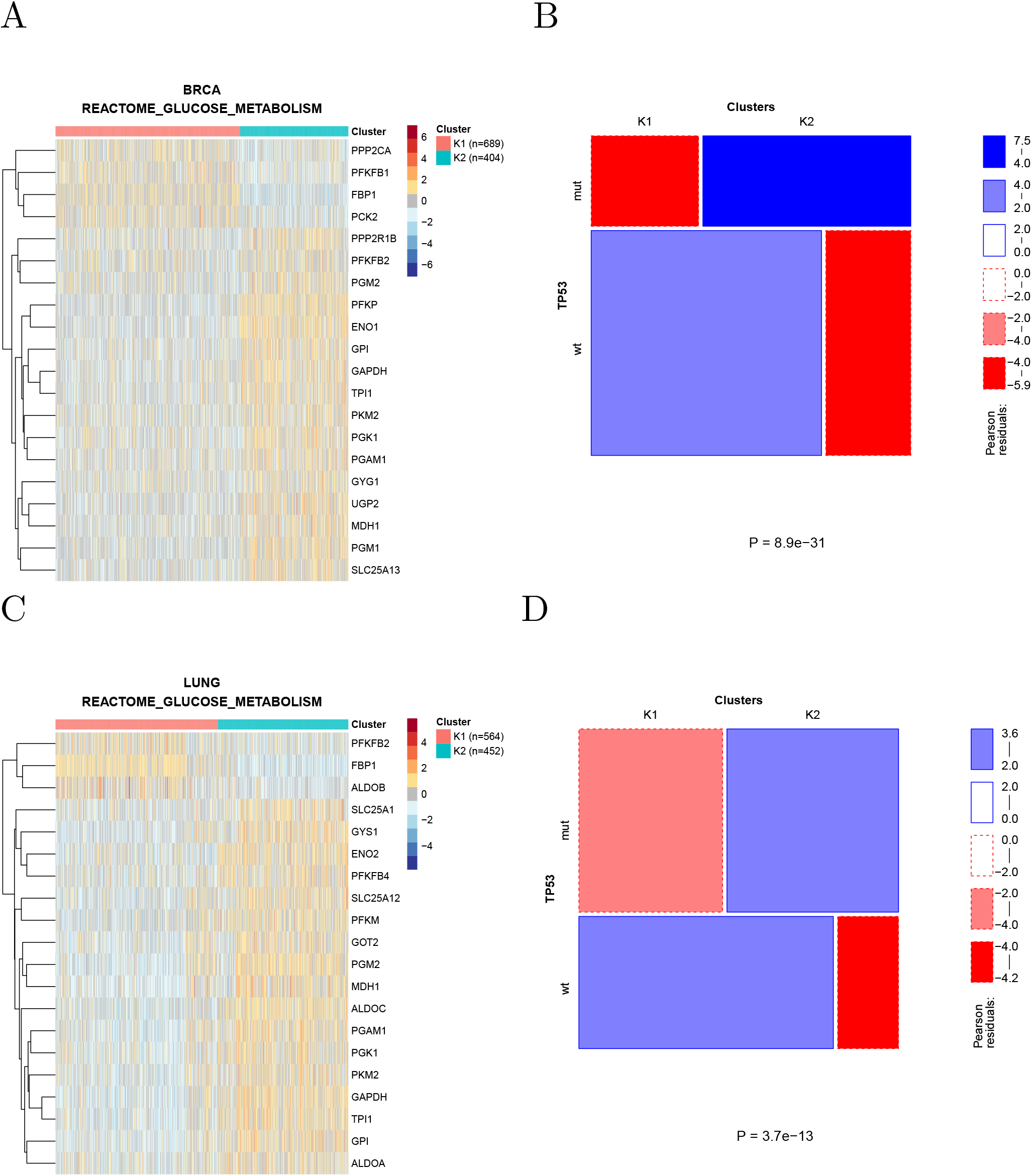
(A) Clustering of breast cancer samples using the genes associated to glucose metabolism according to REACTOME. Cluster K2 shows upregulation of Warburg effect genes such as GAPDH, PGK1, and PKM2 (B) The prevalence of TP53-mutated samples is significantly higher in cluster K2. (C, D) Same for lung cancer samples

### Pyruvate metabolism in thyroid cancer

The clustering of thyroid cancer samples according to the genes associated by KEGG to pyruvate metabolism is shown in Fig. 7A. These 2 clusters show many significant differences in both clinical and molecular parameters. In particular (Fig. 7B,C), the prevalence of BRAF and NRAS-mutated samples is strikingly different in the two clusters, with *all* NRAS-mutated samples found in cluster 2 and the large majority of BRAF-mutated samples found in cluster 1. The two clusters also strongly differ in the respective prevalence of histological types.

**Figure 7:**
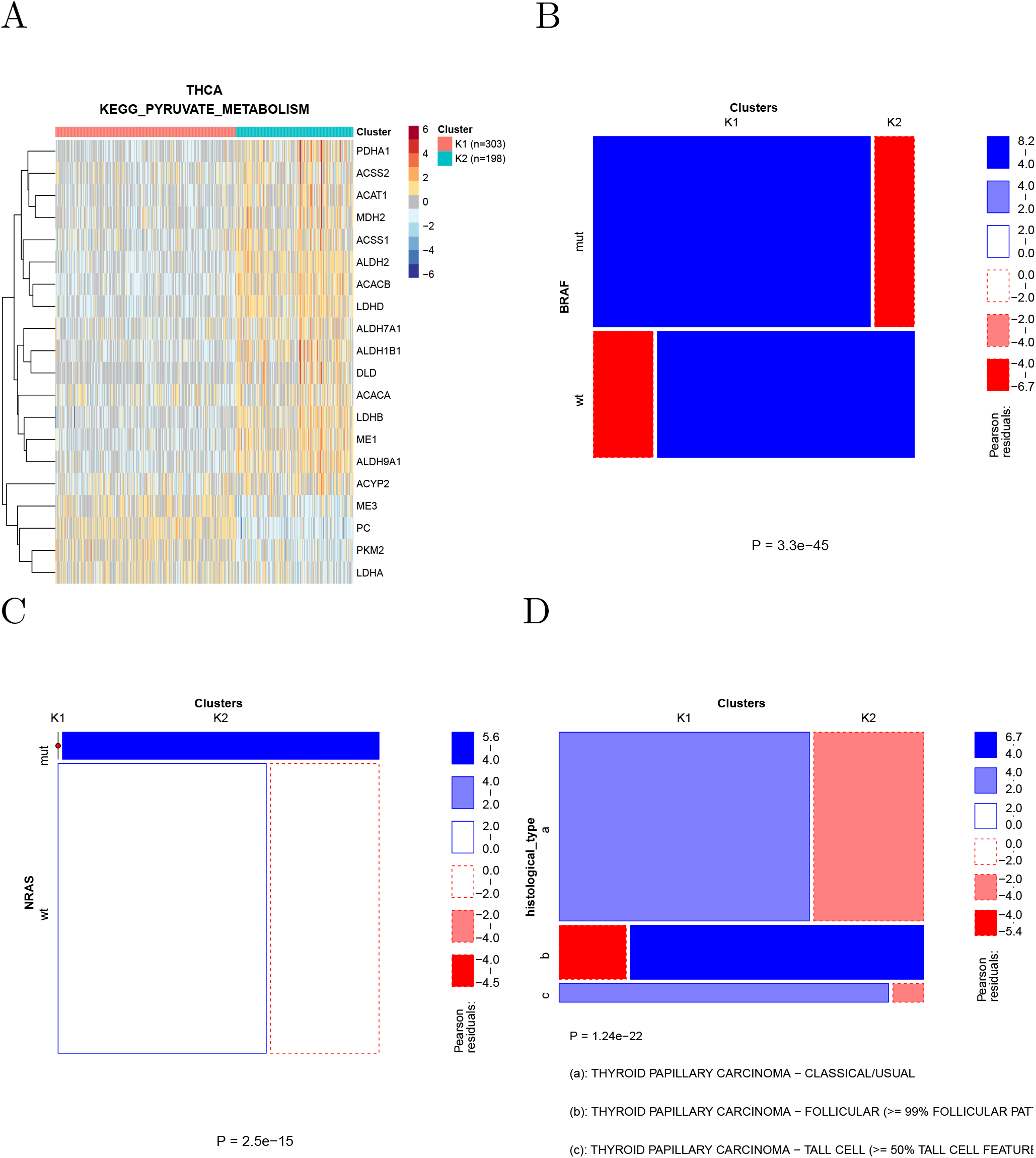
(A) Clustering of thyroid cancer samples using the genes associated by KEGG to pyruvate metabolism. (B) Most BRAF-mutated samples fall into cluster 1 (C) All NRAS-mutated samples fall into cluster 2. (D) The two clusters significantly differ in the prevalence of histological types.

### Fatty acid metabolism in thymoma

Clustering of thymoma samples using the genes associated to fatty acid metabolism divides the samples in two clusters (Fig. 8A) which largely overlap the known histological types (Fig. 8B) and differ in the prevalence of a recurrent GTF2I mutation [12].

**Figure 8:**
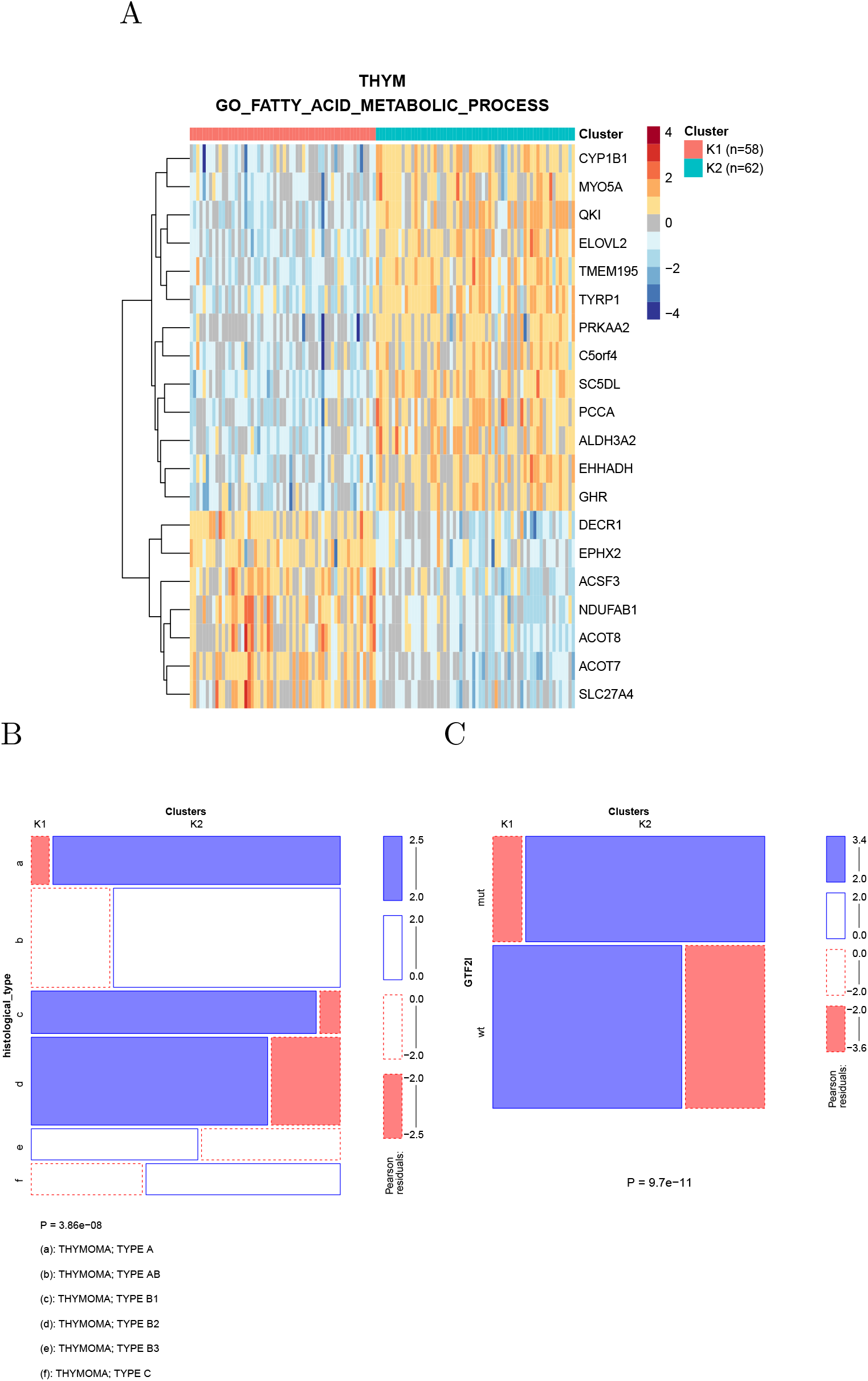
(A) Clustering of thymoma samples using the genes associated by KEGG to fatty acid metabolism. (B) The clusters significantly overlap the known histological types (C) A specific GTF2I mutation signifcantly segregates between the two clusters

### Analysis of cancer cell lines

We performed the same analysis on the cancer cell lines included in the cancer cell line encyclopedia [13]. These cell lines are classified in terms of TCGA tumor types (22 types are represented by a total of 723 CCLE cell lines). Within each tumor type, cell lines were clustered based on their transcriptomic data, using the expression of the genes in the same metabolic gene sets used to cluster TCGA samples. The clusters were then correlated with molecular and phenotypic data, including copy number alterations, specific point mutations, gene-level mutations, protein expression, microRNA expression, and drug response data (IC50 values).

Since the number of cell lines in the CCLE is about one order of magnitude smaller that the number of TCGA, samples, the power to detect significant associations is also much smaller. Nevertheless, we found 485 significant associations at a Bonferroni-corrected significance level of 0.05. For example, KEAP1 mutations are associated with xenobiotic metabolism in both TCGA samples and CCLE cell lines (Fig. 9), in agreement with the known role of the NRF2/KEAP1 pathway in the regulation of xenobiotic response [14]. Note that KEAP1 mutations appear to be associated to the activation of the pathway, as observed in [15].

**Figure 9:**
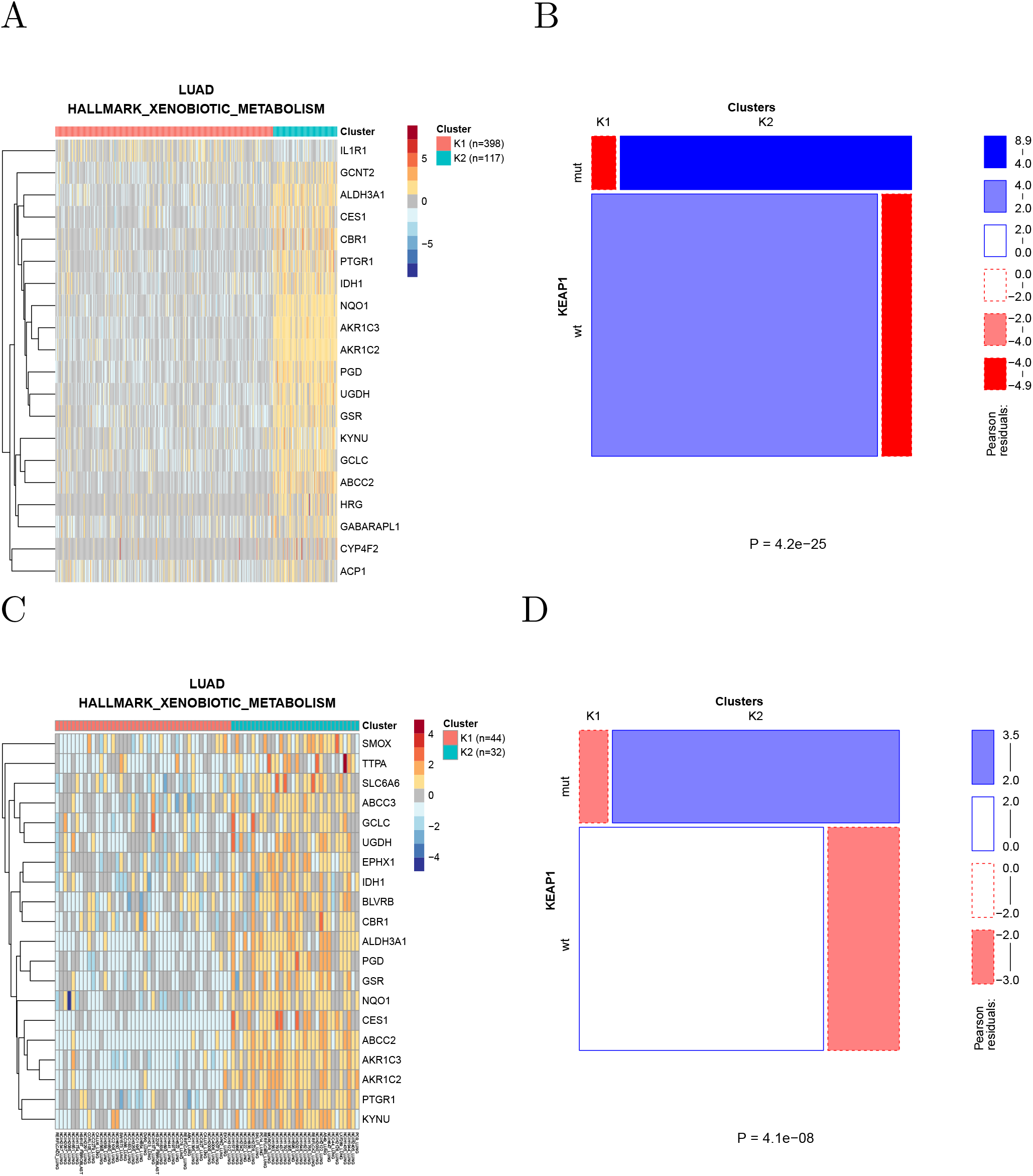
(A) Clustering of lung adenocarcinoma samples using the genes in the xenobiotic metabolism hallmark gene set. (B) Mutations of KEAP1 are strongly enriched in the K2 cluster where the pathway appears to be activated. (C) Clustering of lung adenocarcinoma cell lines usineg the same gene set. (D) Also in cell lines mutations are enriched in the cluster showing pathway activation.

## Discussion

In this work we systematically analyzed cancer metabolic subtypes, defined through clustering of patients using the transcription profiles of metabolic gene sets. These subtypes are highly specific of each individual metabolism used to define the gene sets, and show a myriad of significant associations with both molecular and phenotypic features of the tumors.

We present these associations as a resource for the community, available from metaminer.unito.it which we believe will be useful in two related but distinct ways. One one hand, our results highlight the key role of metabolic profiles in classifying individual patients in a way that is potentially useful to devise precision treatments. This is particularly compelling for tumors for which no established classification exists, but in which metabolic subtypes strongly associated with phenotypic characteristics can be found.

On the other hand, our results can help formulating hypotheses on the mechanisms underlying the associations between metabolic subtypes on one hand and molecular and phenotypic features on the other, which can then be experimentally tested in cell lines or organoids. For example, while both miR-222 (see e.g. [16] for a recent review) and inositol phosphate metabolism [17,18] have been often associated to cancer progression, their association, which recurs in a large number of tumor types, has not been previously reported to the best of our knowledge, and deserves a mechanistic investigation.

Indeed the correlative nature of our results is the most obvious limitation of our results, and of all results obtained by the retrospective analysis of primary tumors.

## Methods

### Data

#### TCGA data

All TCGA data (release 2016_02_28) were obtained from the Broad TCGA GDAC site (https://gdac.broadinstitute.org/), by means of firehose_get, version: 0.4.1. The gene expression data clustered to generate the metabolic subtypes were the normalized gene-level TPMs obtained with RSEM.

Copy number data were obtained from the segmented data using the cghMCR [19], DNAcopy [20], and CNTools [21] Bioconductor packages. The cghMCR package allows the calculation of segment gain or loss (SGOL) from segmented data, by means of a modified version of the GISTIC algorithm. The *segment* function of the DNAcopy package is used to segment the normalized data so that chromosome regions with the same copy number have the same segment mean values. Then, using the *getRS* function from CNTools, the data returned by segment are organized in a matrix format. Finally the *SGOL* function of cghMCR is used to compute gene-level SGOL scores for genes by calculating the sum (parameter method) of all the positive and negative values, respectively above and below a set threshold (0.5, −0.5). Finally, SGOL scores for all genes included in each Broad Institute positional gene set were averaged to obtain the SGOL score of each region.

Genome-wide DNA methylation was obtained from the gene-level summarized Human Methylome 450k data, as provided by GDAC, then summed over all genes to associate a global DNA methylation value to each sample. miRNA expression was obtained from the Illumina HiSeq data as deposited in GDAC. Specific point mutations for each sample were obtained from the “mutation packager oncotated calls” provided by GDAC, considering only non-synonymous mutations. For gene-level mutations, a sample was considered as mutated in a gene if it carried one or more specific point mutations associated to the gene.

Protein expression data (RPPA) were obtained from the corresponding files annotated with gene names from the GDAC site. Survival and other clinical data were obtained from the “Tier 1 clinical pick” files available from GDAC.

#### CCLE data

Gene expression data of 723 cell lines associated to one of the TCGA tumor types were obtained, as gene-level TPMs computed with RSEM, from the CCLE web site. We also retrieved from the CCLE web site the following data to be correlated to metabolic subtypes:

- gene-level copy number data: the absolute score provided by CCLE was processed with the same steps used for the SGOL score reported for the TCGA data to obtain region-based copy number data
- miRNA expression data (which were log-transformed for display in the figures)
- somatic mutation data (also here we considered only non-synonymous mutations)
- IC50 values for the available drugs
- metabolite levels

#### Metabolic gene sets

Gene sets were obtained from the c2.KEGG, c2.REACTOME, c5.BP and Hallmark MSigDB v5.2 collections. Metabolic gene sets were defined as those whose name matched the string “metabol” but not the string “regul” (to include only gene sets directly involved in metabolic pathways rather than in their regulation). In this way we obtained 345 metabolic gene sets (41, 35, 265, and 4 for the c2.KEGG, c2.REACTOME, c5.BP, and Hallmark collections, respectively).

### Generation of metabolic clusters

For each tumor type and each metabolic gene set, the patients were divided into metabolic subtypes using partition around medoid (PAM) clustering [9] on the Spearman rank correlation coefficient-based distance matrix obtained from the expression of the genes in the gene set. The number of clusters (the maximum being set at 10) generated by each metabolic gene set was based on optimizing the average silhouette width. The normalized mutual information between two clusterings of the same patients based on different gene sets was computed with the R package *infotheo* [22], using the entropy of the empirical probability distribution and normalizing to the geometric mean of the entropies. Genes differentially expressed among clusters were identified using a Kruskal-Wallis test: the top 20 genes by P-value are shown in the heatmaps.

### Statistical tests of association

The statistical tests used for establishing the association between each clinical/molecular feature and metabolic subtypes depend on the type of variable describing the feature. The metabolic subtype is always a categorical variable indicating the cluster membership of each patient.

- For continuous variables (microRNA expression, DNA methylation, CNA SGOL scores) we used Mann-Whitney U test (Kruskal-Wallis for *k* > 2). In the case of microRNA expression, given the large number of microRNAs with expression equal to zero in each sample, the expression data were rank-transformed with random resolution of ties before the statistical test.
- For categorical variables (e.g. presence of a given mutation) we used Fisher’s exact test (*χ*^2^ test for tables larger than 2 × 2, removing levels for which the expected count was less than 5)
- For survival (RFS or OS) we used the log-rank test. The test was not considered valid when the expected number of events in any metabolic subtype was less than 5

#### Multiple testing

Bonferroni correction was applied separately to each variable type but to all tumor types together. Thus, for example, the Bonferroni correction for the association with miRNA expression takes into account all tests of association between all miRNAs and all metabolic gene sets across all tumor types. We chose to perform the correction separately for the feature types because these contain very different numbers of variables (hundreds of variables for miRNA expression and a single one for overall survival): Therefore an overall Bonferroni correction would unduly penalize the variable types containing few variables.

## Supporting information

Supplementary data 1: TCGA clusters

Supplementary data 2: TCGA associations

Supplementary data 3: CCLE clusters

Supplementary data 4: CCLE associations

## Data availablilty

The results of the analysis can be browsed interactively at https://metaminer.unito.it/.

## Supplementary material

- Supplementary data 1: clustering of TCGA patients into metabolic subtypes.
- Supplementary data 2: associations between metabolic subtypes and molecular/phenotypic features in TCGA.
- Supplementary data 3: clustering of CCLE cell lines into metabolic subtypes.
- Supplementary data 4: associations between metabolic subtypes and molecular/phenotypic features in CCLE.

## Acknowledgements

The results of this analysis are in whole or part based upon data generated by the TCGA Research Network: https://www.cancer.gov/tcga. We would like to thank Davide Cittaro for insightful comments and discussions; and Alessandro Greganti, Ivan Molineris, and Sergio Rabellino for IT support.

